# Heterogeneity of hypothalamic Pro-opiomelanocortin-expressing neurons revealed by single-cell RNA sequencing

**DOI:** 10.1101/103408

**Authors:** Brian Y. H. Lam, Irene Cimino, Joseph Polex-Wolf, Sara Nicole Kohnke, Debra Rimmington, Valentine Iyemere, Nicholas Heeley, Chiara Cossetti, Reiner Schulte, Luis R. Saraiva, Darren W. Logan, Clemence Blouet, Stephen O’Rahilly, Anthony P. Coll, Giles S. H. Yeo

**Author notes:** Corresponding Author Giles SH Yeo (;) University of Cambridge Metabolic Research Laboratories, MRC Metabolic Diseases Unit, Wellcome Trust-MRC Institute of Metabolic Science, Box 289, Addenbrooke’s Hospital, Cambridge CB2 0QQ, UK. **Accession Numbers:** GSE92707.

## Abstract

Arcuate proopiomelanocortin (POMC) neurons are critical nodes in the control of body weight. Often characterised simply as direct targets for leptin, recent data suggest a more complex architecture. Using single cell RNA sequencing, we have generated an atlas of gene expression in murine POMC neurons. Of 163 neurons, 118 expressed high levels of *Pomc* with little/no *Agrp* expression and were considered “canonical” POMC neurons (P^+^). The other 45/163 expressed low levels of *Pomc* and high levels of *Agrp* (A^+^P_+_). Unbiased clustering analysis of P^+^ neurons revealed four different classes, each with distinct cell surface receptor gene expression profiles. Further, only 12% (14/118) of P^+^ neurons expressed the leptin receptor (*Lepr*) compared with 58% (26/45) of A^+^P_+_ neurons. In contrast, the insulin receptor (*Insr*) was expressed at similar frequency on P^+^ and A^+^P_+_ neurons (64% and 55%, respectively). These data reveal arcuate POMC neurons to be a highly heterogeneous population.

## Introduction

The leptin-melanocortin pathway plays a key role in the control of food intake and body weight, with genetic disruption of multiple components of the pathway resulting in severe, early-onset obesity in both humans and mice (reviewed by (Yeo and Heisler, 2012)).

In the brain, POMC expression is localized to neurons in the arcuate nucleus (ARC) of the hypothalamus and in the nucleus of the solitary tract (NTS) (Cone, 2005). POMC is a pro-peptide that is processed to a range of bioactive peptide products including α and β-MSH (Bertagna, 1994; Castro and Morrison, 1997), both of which act on melanocortin receptors 3 and 4, which are widely expressed in the central nervous system. The role of MC4R in mediating the effects of POMC on energy balance is well established, whereas the role of MC3R is less clear (Cone, 2005). In the ARC, anorexigenic POMC neurons exist alongside orexigenic neurons that express the endogenous melanocortin antagonist agouti-related peptide (AgRP) (Fan et al., 1997; Ollmann et al., 1997). The canonical view of the melanocortin pathway is one where leptin, which is produced from white adipose tissue to reflect fat mass, signals to leptin receptors on the POMC and AgRP neurons, increasing the activity of POMC neurons and reducing the activity of AgRP expressing neurons (Cone, 2005; Yeo and Heisler, 2012).

However, there is a body of evidence that indicate this view of the leptin melanocortin pathway to be an over-simplistic representation of the relevant circuitry. Woods and Stock, using FOS immunoreactivity of the immediate early gene *c-fos* as a marker of neuronal activation, reported that while neurons in the arcuate nucleus of leptin deficient *ob/ob* mice were activated by peripheral leptin administration, the area of the hypothalamus most strongly activated was the paraventricular nucleus (Woods and Stock, 1996). Later, it became clear that not all POMC neurons express the receptors for leptin or insulin, with whole-cell patch-clamp electrophysiology indicating that a subpopulation are depolarized by leptin, while a separate and distinct population is hyperpolarized by insulin (Williams et al., 2010). POMC neurons also vary in their expression of the neurotransmitters glutamate, Ƴaminobutyric acid (GABA) (Hentges et al., 2009; Jarvie and Hentges, 2012), and acetylcholine (Meister et al., 2006) and their downstream projections. For instance, 40% of POMC neurons appear to be GABAergic and hence inhibitory (Hentges et al., 2009). In addition, only about 40% of POMC neurons express the serotonin 2C receptor (5HT_2C_-R), the target for the weight-loss drug Lorcaserin, through which it is thought to mediate much of its actions on satiety (Burke et al., 2016; Heisler et al., 2002). Finally, mice with a targeted deletion of the leptin receptor specifically from POMC neurons display a surprisingly mild phenotype (Balthasar et al., 2004), as compared to complete *Pomc* null mice (Challis et al., 2004; Yaswen et al., 1999) and *db/db* mice homozygous for a *Lepr* mutation (Chen et al., 1996).

A recent paper by Henry et al reported the gene expression profiles of pooled AGRP and POMC neurons from fed and food deprived mice, and found that the global pattern of gene expression in response to an acute energy deficit are far more dramatic in AGRP expressing than POMC expressing neurons (Henry et al., 2015). However, because of its use of pooled cells, the transcriptomic profiles of individual neurons, and hence their variability in gene expression, was not examined.

We now report the results of single cell RNA sequencing in isolated POMC expressing neurons of the arcuate nucleus which has revealed a previously unappreciated degree of heterogeneity within this anatomically highly localised population.

## Results

## RNA-sequencing of POMC neurons

Using a POMC-eGFP mouse (Cowley et al., 2001), we successfully sequenced the transcriptome of 163 GFP positive (MBH Pos) and 7 GFP negative cells (MBH Neg) from the hypothalamic tissue encompassing the arcuate nucleus (**Fig 1A**), and 24 cells from the pituitary (Pit) as positive controls known to express high levels of POMC (GEO Acc# GSE92707). When we visualized the transcriptomic data using t-SNE dimension reduction analysis (Maaten and Hinton, 2008), the pituitary and hypothalamic cells segregated into two discrete clusters, with the hypothalamic neurons further segregating into GFP positive and GFP negative cells (**Fig 1B**).

All cells sequenced were of similar size, as measured by degree of forward scatter on the cell-sorter, and expressed both *Actb* and *Gapdh* (**Fig 1C**). All the hypothalamic cells expressed the neuronal marker *Ncam*, but because the pituitary cells contained both neuronal derived posterior and the ectodermally derived anterior, there was a larger heterogeneity of *Ncam* expression in the pituitary cells (**Fig 1C**). In contrast, the GFP signal, which reflects the degree of *Pomc* expression, was consistently high with very little heterogeneity in the pituitary POMC cells, whereas hypothalamic GFP neurons show a much larger variation in signal. *Cartpt*, which is canonically expressed together with *Pomc* in the same neurons, also showed variable expression in hypothalamic GFP neurons and was not detected in the pituitary (**Fig 1C**). POMC is processed by the proprotein convertases, PC1 (PCSK1) and PC2 (PCSK2). Using the detection cut-off ≥5 linear RPM, most cells (156/163) expressed either or both *Pcsk1* and *Pcsk2* (**Fig 1D**). On average, 6882 different transcripts were detected from each individual neuron (≥5 linear reads per million or RPM). However, across all 163 POMC neurons, 12,187 different transcripts were expressed (**Fig 1E**), indicating significant heterogeneity in gene expression repertoire among the neurons.

**Figure 1.**
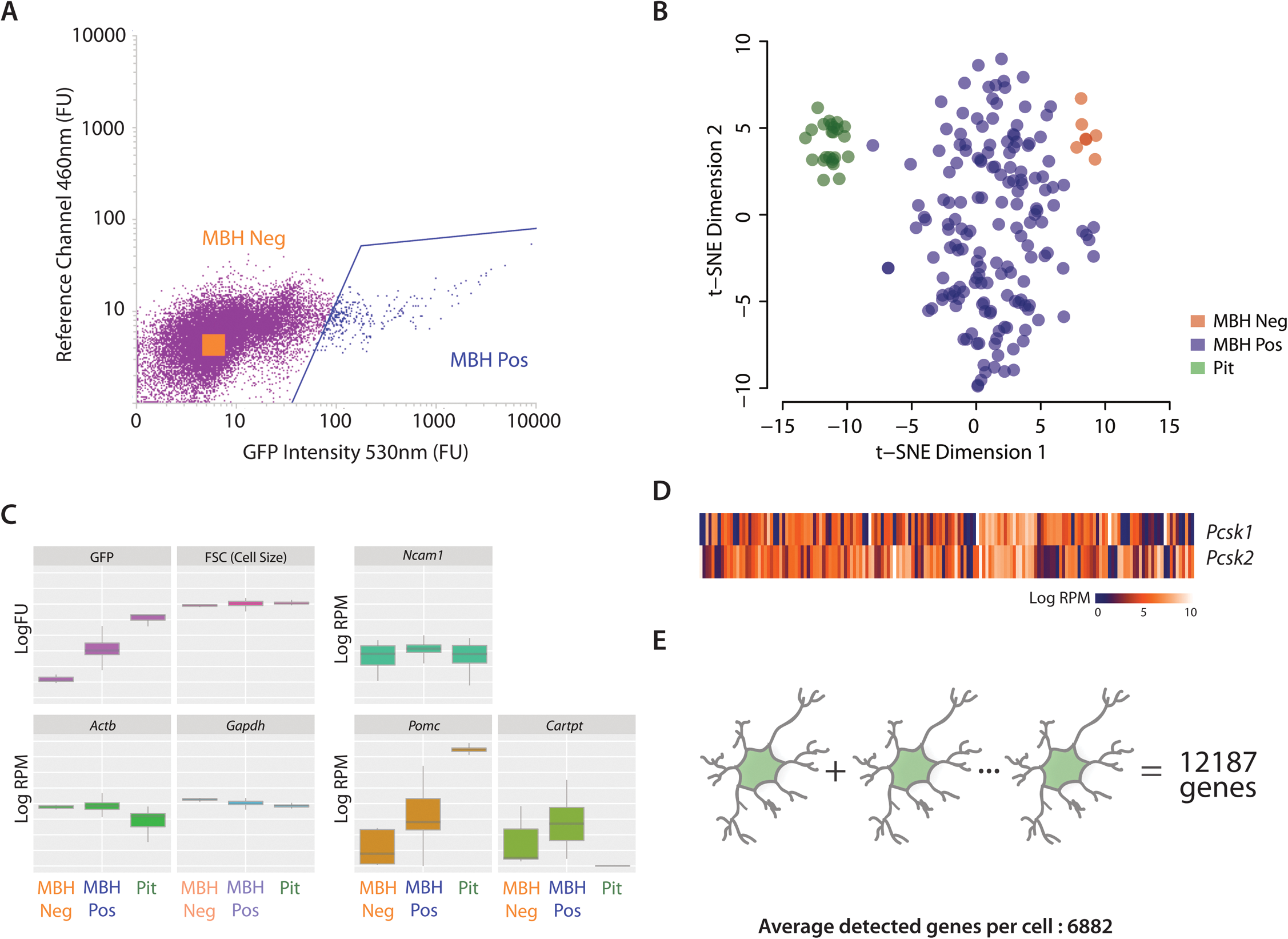
FACS sorting and single-cell RNA sequencing of POMC-eGFP neurons. **(A)** The FACS gating for GFP (530nm) and reference channel (460nm). The GFP positive cells from the mediobasal hypothalamus (MBH Pos) are labelled blue and negative cells (MBH Neg) are collected from the orange square within the negative population (purple). The cells were further gated for nuclear stain (DraQ5), cell size (FSC) and granularity (SSC), data not shown. **(B)** t-SNE was used to show the segregation of cells (MBH Pos and Neg, as well as GFP positive cells collected from the pituitary) based on their transcriptomic profiles. **(C)** Quality checks for FACS showing the GFP intensity and cell size measurement amongst the 3 cell types. We also examined for RNA expression of housekeeping genes *Actb, Gapdh,* neuronal markers *Map2* and *Ncam1*, and also expression of *Pomc and Cartpt*. **(D)** *Pcsk1* and *Pcsk2* are widely expressed in the cells sequenced. **(E)** We detected an average of 6882 genes per cell (≥5 RPM), adding all the genes from all the 163 cells together resulted in a total of 12187 detected genes, indicating heterogeneity in gene expression.

## AgRP/NPY expressing POMC neurons

Unexpectedly, we found that 27% (45/163) of *Pomc*-expressing neurons also expressed high levels of the endogenous melanocortin antagonist *Agrp* (**Fig 2A**). In addition, these neurons expressed a high level of *Npy*, while *Pomc* expression itself was a magnitude (>5 logFC) lower than that seen in POMC neurons not expressing or expressing low levels of *Agrp* (**Fig 2B**). Comparing our data with a published transcriptome of pooled AgRP neurons (Henry et al., 2015), we found a significant similarity in gene expression profiles (R^2^ = 0.71 *p*<0.00001), with high expression, for instance, of the ghrelin receptor *Ghsr* (**Fig 2B**). Analysis of the raw data from Henry et al 2015 also indicated that *Pomc* and *Cartpt* were expressed in the AgRP neurons that they had analysed (Henry et al., 2015), albeit at low levels (**Fig 2D**).

Taken together, these 45 high *Agrp* and low *Pomc* expressing neurons (hereafter referred to as A^+^P_+_ neurons) showed a striking similarity to canonical AgRP/NPY neurons, likely reflecting their developmental origins, with up to 25% of AgRP neurons sharing the same hypothalamic progenitors as POMC neurons (Padilla et al., 2010). We defined the rest of the 118 neurons, which expressed higher levels of *Pomc*, as canonical POMC expressing neurons (P^+^) and conducted our further studies on this population.

**Figure 2.**
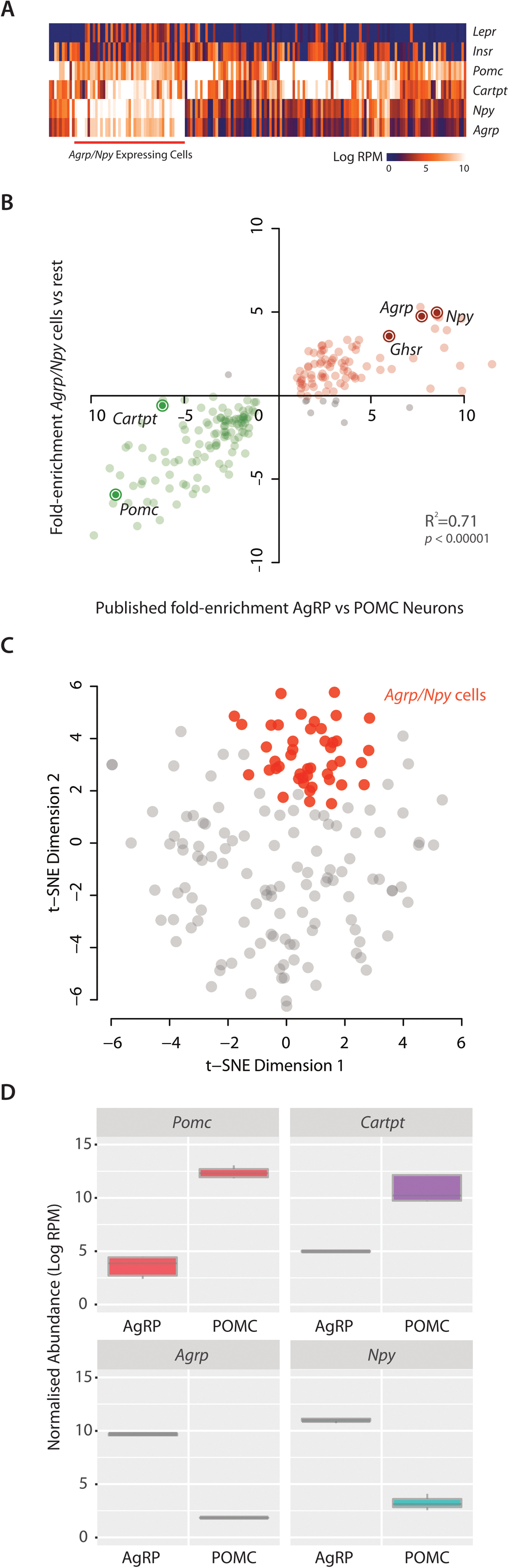
27.6% of POMC cells also express high levels of *Agrp* and *Npy*. **(A)** We detected 45 cells that expressed high levels of *Agrp* and *Npy* (A^+^P_+_), interestingly many of these cells also express *Lepr*. **(B)** Differential expression analysis of the A^+^P_+_ vs rest (P^+^) cells shows similar fold enrichments compared to AgRP vs POMC comparison from (Henry et al., 2015). The expression of the top 200 enriched genes (from the published data) are plotted in the graph. **(C)** The *Agrp/Npy* cells form a strong cluster in a t-SNE plot showing only the 163 MBH Pos cells. **(D)** Expression levels of *Pomc*, *Cartpt*, *Agrp*, and *Npy* in published AgRP and POMC neurons. The AgRP neurons express low but detectable levels of *Pomc* and *Cartpt*, on the other hand *Npy* and *Agrp* are also detectable in POMC neurons.

## Expression of hypothalamic neuroendocrine genes in POMC neurons

We produced a matrix showing the expression distribution of 23 well characterised neuroendocrine genes, all known to play a role in the regulation of energy homeostasis, on the individual POMC (P^+^) neurons (**Fig 3A**). We found, as has been previously reported, that a large percentage of POMC neurons expressed *Mc3r*, *Cnr1*, *Htr2c* and the *Npy-y1*, *y2* and *y5* receptors (Yeo and Heisler, 2012). Of note, the percentage of *Htr2c* expressing neurons (34/118; 29%) was comparable to previous studies showing that 5HT_2C_Rs (the protein product of *Htr2c*) are expressed on approximately 40% of POMC neurons (Burke et al., 2016). We also found that a smaller percentage of neurons, 12/118, expressed *Mc4r* (**Fig 3B**), with 11/118 expressing *Glp1r*. ARC POMC neurons expressing GLP1R are thought to mediate, in part, the weight-loss effects of the GLP1 analogue liraglutide (Secher et al., 2014).

**Figure 3.**
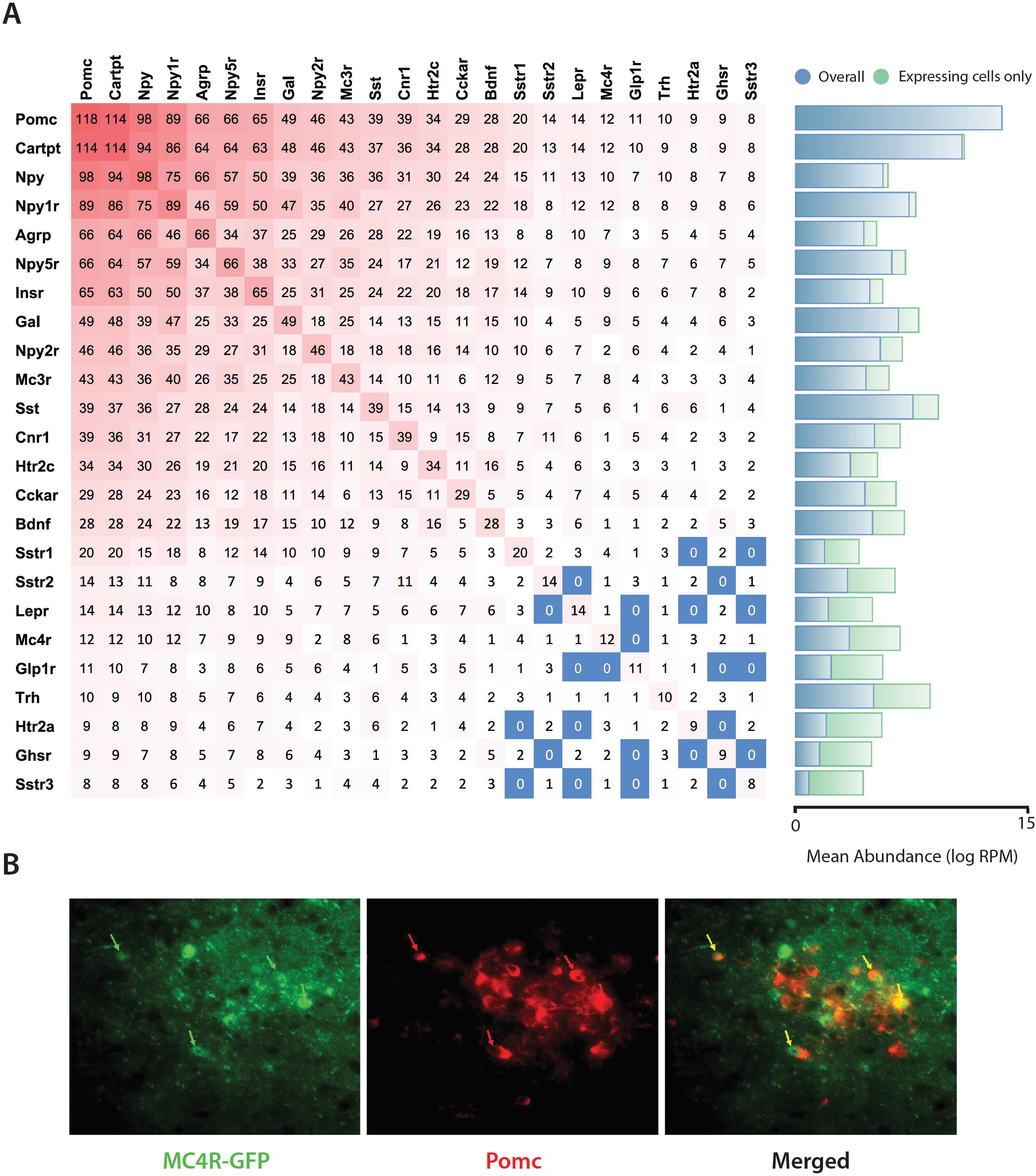
The expression matrix of selected neuroendocrine genes. **(A)** An expression matrix showing the number of cells expressing ≥2.32 log RPM of a selection of neuroendocrine genes, the *Agrp/Npy* cells were excluded from the matrix. One can look up to coexpression of up to 2 genes by looking at the intersections between rows and columns, e.g. there are 43 *Mc3r* expressing cells (9^th^ row) and 25 of them also expressed *Insr* (7^th^ column). The average abundance of gene expression is also shown in blue bars (average of all cells in the cohort) and green (expressing cells only) **(B)** We found 12 cells expressing *Mc4r* within our cohort and this was confirmed by immunostaining for POMC the arcuate nucleus of MC4R-GFP mice.

While we excluded the 45 A^+^P_+_ neurons from this matrix, a large number of the remaining P^+^ neurons still expressed low levels of *Agrp* (66/118) and *Npy* (98/118). Our own data (**Fig 3A**) and re-analysis of the raw data from previously published work indicated that the converse was also likely to be true, with a percentage ‘bona fide’ AgRP neurons also expressing low levels of *Pomc* (**Fig 2D**),

## Unbiased clustering reveals four ‘classes’ of POMC neurons

In order to determine whether P^+^ neurons segregate into distinct classes, we first adjusted for batch effects and then performed unbiased clustering of the 118 cells based on the consensus matrix of Euclidean, Pearson and Spearman correlations - using the ‘SC3’ R package (Kiselev et al., 2016). We obtained 4 main clusters (**Fig 4A**), which could also be visualized using t-SNE (Maaten and Hinton, 2008) (**Fig 4B**). Cluster 1 was the largest with 69 cells, followed by cluster 4 (27) and cluster 2 (15). Cluster 3 had the least number of cells (7). Cluster 2 showed resemblance to cluster 1 (**Fig 4A, B**).

We identified 2773 candidate genes that were differentially expressed among the 4 clusters, with a false discovery rate (FDR) cut-off of 5%, that drove the segregation of cells into the different groups (**Fig 4C**). Interestingly, there was over-representation of membrane (530) and secretory genes (191) within the driver gene list (**Fig 4C, D**). These membrane genes encode ion channels, G-protein coupled receptors (GPCRs) and other transmembrane receptors, and transporters (**Fig 4D**).

**Figure 4.**
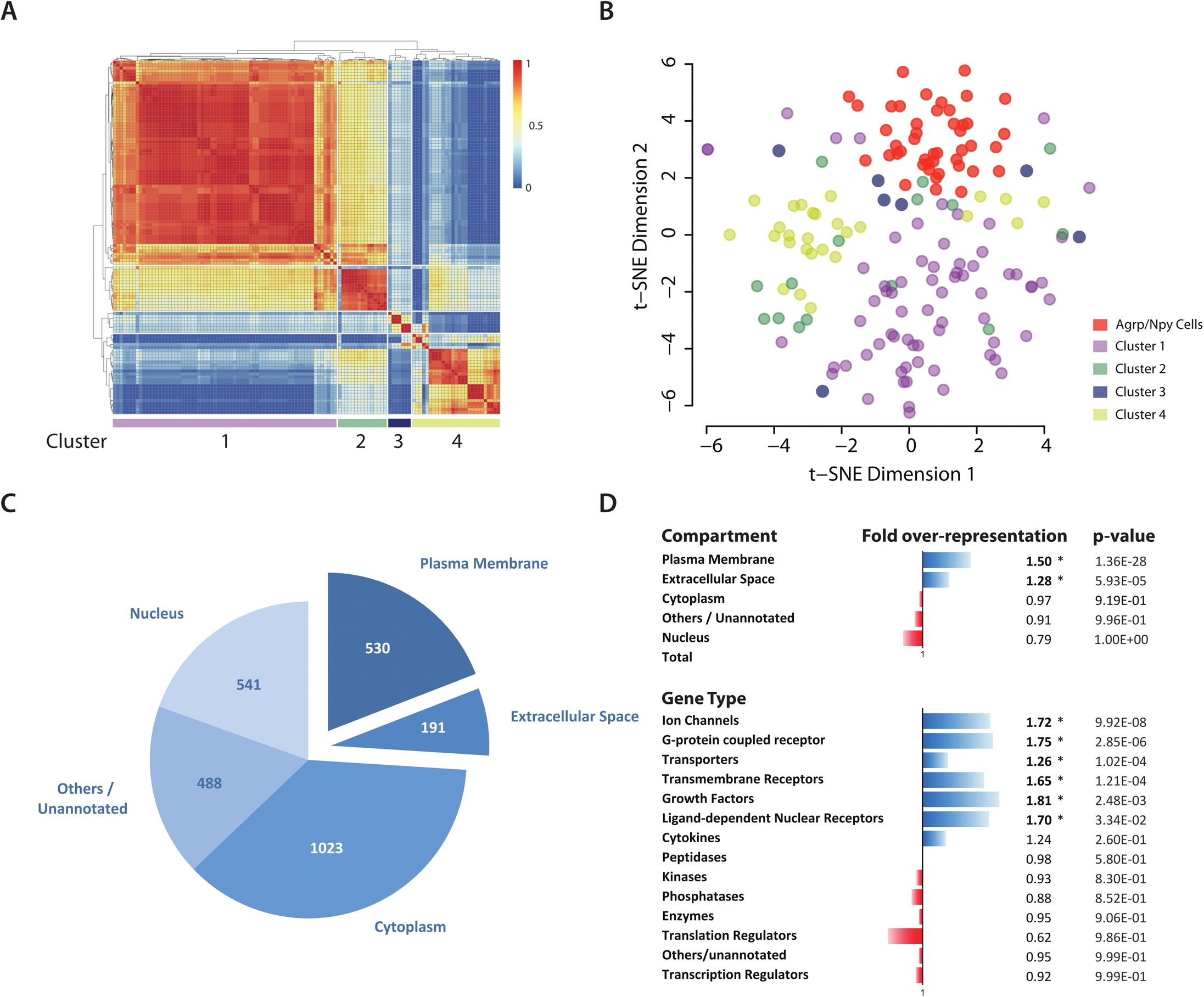
Clustering analysis and identification of driver genes for heterogeneity. **(A)** Clustering analysis was performed using the ‘SC3’ package in order to identify clusters of cells that show similarity in their transcriptomic profiles. Using k-means clustering, we found 4 clusters with an average silhouette width of 0.64. **(B)** A t-SNE plot showing the 4 clusters from the analysis. **(C)** Using the clustering results we identified 2773 candidate genes that drove heterogeneity. **(D)** There are over-representations of genes from the plasma membrane (1.50-fold) and extracellular space (1.28-fold) within the list of driver gene candidates. This enrichment is mainly driven by overrepresentations of plasma membrane protein encoding genes including ion channels, G-protein coupled receptors, transporters and other transmembrane receptors. There was also a 1.8-fold enrichment of growth factors in the driver gene list.

**Figure 5A** lists the top 20 genes, ranked by differential expression that drove the clustering of the four groups (with the exception of group 2, which was a small group and only has 3 significantly differentially expressed genes). Shown are the expression level for each of the genes and how unique each gene was to each group. Additionally, **Table 1** contained genes that are not in the top 20, encoding selected extracellular secreted hormones, GPCRs and ion channels, indicating which groups had the highest expression of any given gene. For example, while the neurons in cluster 4 expressed the highest levels of *Pomc* and *Cartpt*, both genes were, as expected, also expressed in Clusters 1-3. Cluster 4 neurons however were characterised by expression of genes encoding transthyretin (*Ttr*), the cholesystekinin A receptor (*Cckar*), neuromedin U receptor 2 (*Nmur2*) and nascent helix loop helix 2 (*Nhlh2*) (**Fig 5A**), as well as the GPCRs corticosterone releasing hormone receptor 1 (*Crhr1*), hypocretin receptor 1 (*Hcrtr1*), neuropeptide Y 2-receptor (*Npy2r*) and prolactin releasing hormone receptor (*Prlhr*) (**Table 1**) at far higher levels than in the other clusters. Cluster 1, in contrast, was characterized by high expression of the serotonin-2a receptor (*Htr2a*), the melanocortin-3 receptor (*Mc3r* and a large number of ion channels (**Table 1**).

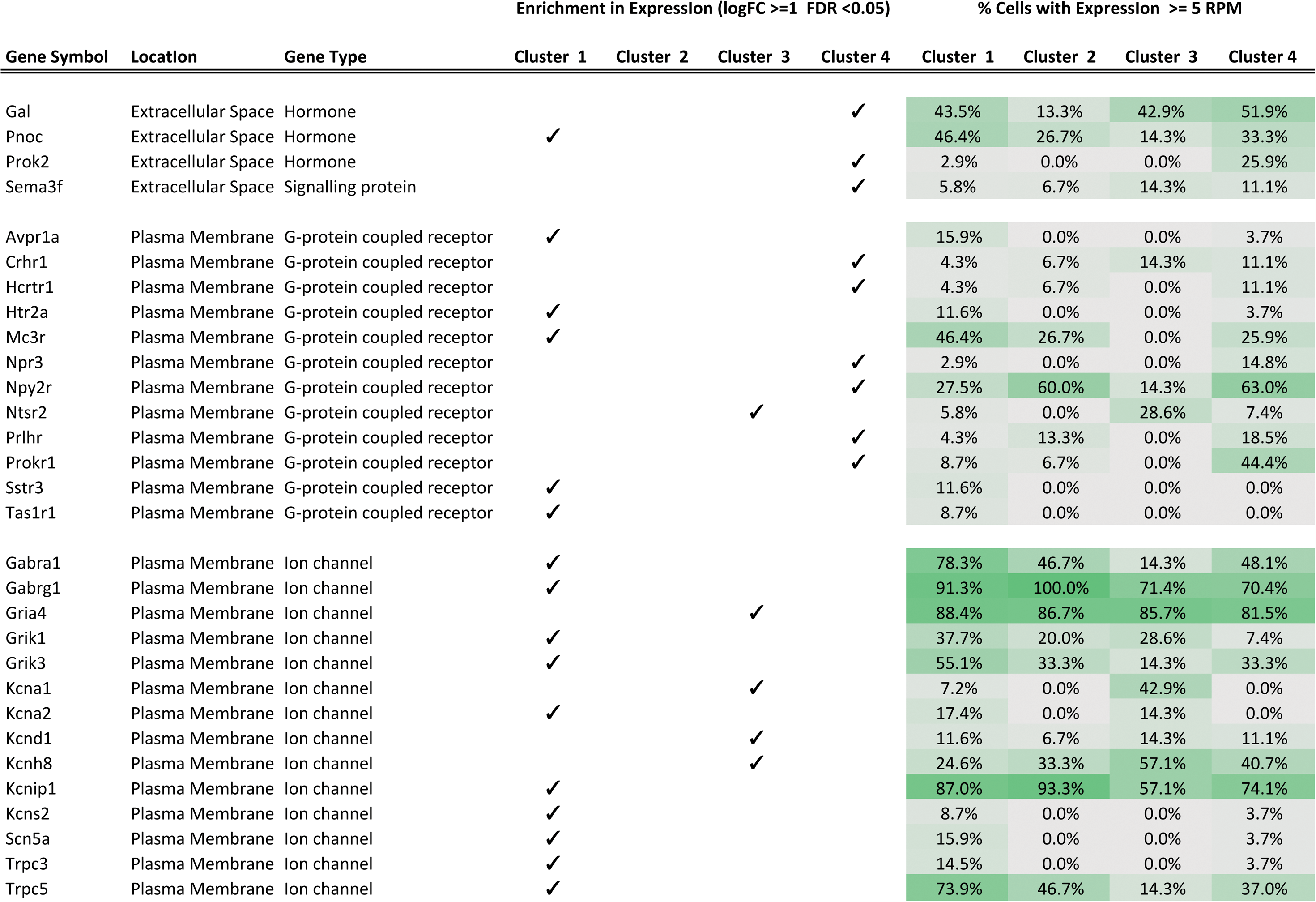

Thus, no single given ‘driver’ gene is responsible for these clusters, but rather the combined expression of a repertoire of genes drove the segregation. The entire repertoires of genes for each cluster are listed in supplementary table 1.

These clusters are, of course, not fixed. For example, **Figure 5B** shows the t-SNE plot with the cluster 4 neurons highlighted and dividing clearly into two groups; one larger group of 22 and a smaller one of 5 neurons. Comparing the transcriptomes of the two groups, we could identify differentially expressed genes that defined the two groups (**Fig 5C**). The four clusters, as we report here, were achieved by using a specific statistical threshold. If we raised this threshold, and therefore statistical rigour, then all clusters, could be further segregated and, if taken to its extreme, could segregate into the original 163 individual neurons sequenced.

**Figure 5.**
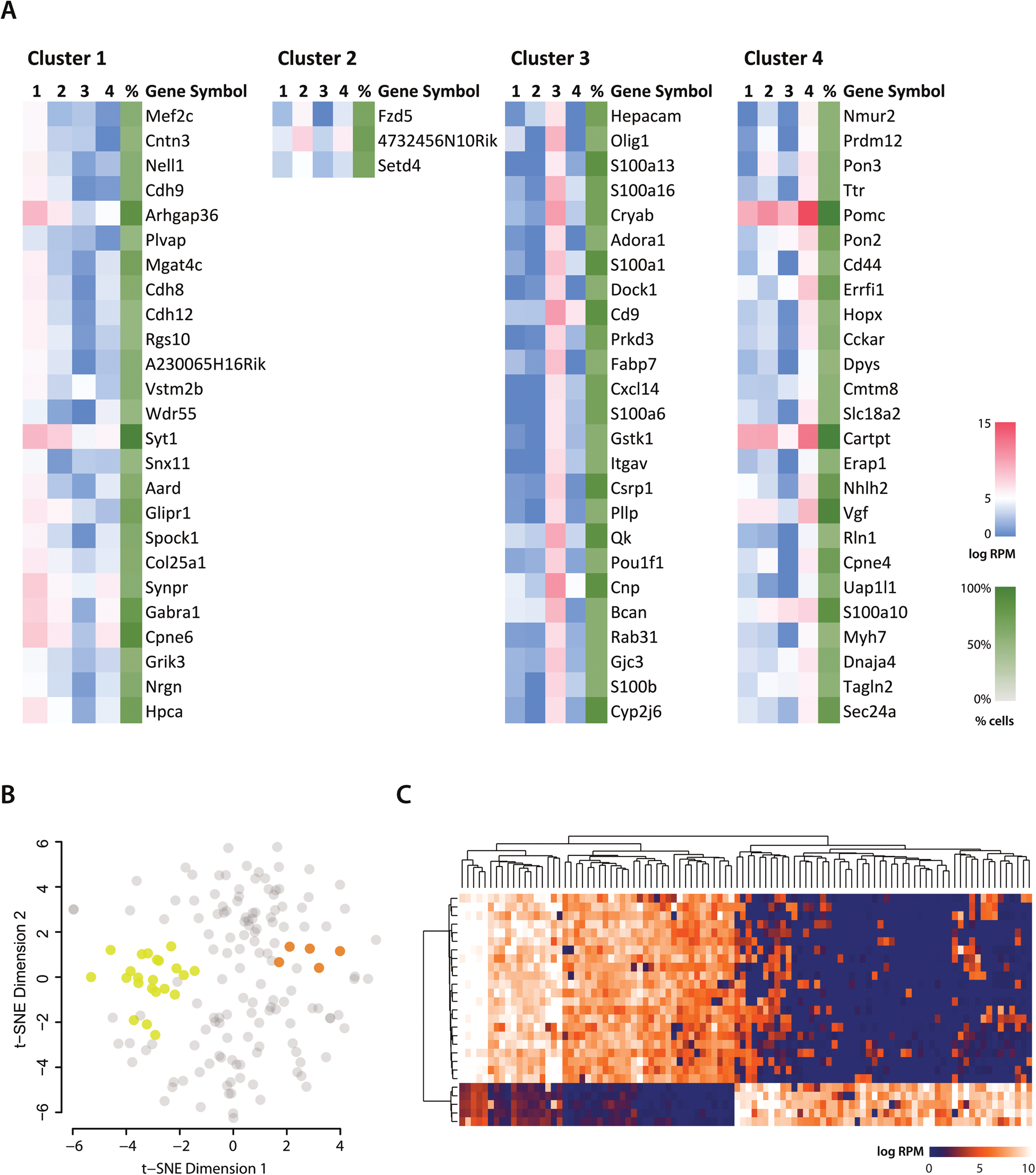
Top candidate driver genes for the resulting clusters. **(A)** The top 25 genes ‘markers’ were ranked by maximum fold difference and a differential expression FDR cutoff of 0.05, and the lists for each of the clusters are shown. The number of cells expressing the gene is also included in the rightmost column (green). *Pomc* and *Cartpt* are expressed in all 4 clusters but the expression of the two genes are highest in Cluster 4 cells (> 1logFC from cluster 2 which is second highest). **(B)** The clusters can be further subdivided, e.g. cluster 4 showed here. **(C)** Top differential expressed genes between the two subgroups of cluster 4 cells. A larger version of the heatmap is available in the supplementary material.

## Leptin receptor and insulin receptor expressing POMC neurons

Next we examined the two most well studied types of POMC neurons, those expressing the leptin (*Lepr*) and insulin receptors (*Insr*). Looking at all 163 neurons, our sequencing data showed that 40/163 (25%) neurons expressed the leptin receptor and 94/163 (57%) expressed the insulin receptor. 11 (7%) express only *Lepr* and *not Insr*, 65 (40%) express only *Insr* and *not Lepr*, while 29 (18%) express both receptors. 58 (36%) do not express either of the receptors (**Fig 6A**).

When we split the 163 cells into the high *Agrp* expressing A^+^P_+_ neurons and high *Pomc* expressing P^+^ neurons, and then looked at the distribution of *Lepr* in these two groups separately, we saw that 26/45 (58%) of A^+^P_+_ neurons expressed *Lepr* as compared to only 14/118 (12%) of P^+^ neurons, almost eight fold more. In contrast, the *Insr* is expressed in 29/45 (64%) of A^+^P_+_ neurons, which is comparable to its expression in 65/118 (55%) of P^+^ neurons.

To determine if we had introduced any selection bias during the flow sorting, we sought to define the number *Lepr* and *Insr* expressing POMC neurons there were in the intact mouse hypothalamus. Attempts to use immunocytochemistry to detect either the leptin or the insulin receptor on whole brain slices proved to be technically unreliable (data not shown). We therefore measured levels of Stat3 phosphorylation (pStat3) after leptin administration and levels of Akt phosphorylation (pAkt) after insulin administration using immunohistochemistry, as a proxy for expression of the respective receptors. We performed these experiments on POMC-GFP mice, allowing us to co-localize pStat3 or pAkt immunofluorescence in neurons expressing GFP (**Fig 6B, C**).

After leptin administration, pStat3 positive cells were largely concentrated to the base of the arcuate nucleus surrounding the median eminence (**Fig 6B**). Cell counting across multiple brain slices from three mice indicated that 23.8% of POMC-GFP neurons were positive for pStat3 in response to leptin, which was consistent with our RNAseq data (**Fig 6C**). Insulin administration however resulted in a different pattern of activation, with pAkt immunofluorescence evident more broadly through the arcuate nucleus (**Fig 6B**). We found that 41.1% of POMC-GFP neurons were positive for pAkt in response to insulin, which was not significantly different (*p*=0.357) to the percentage of GFP neurons expressing *Insr* as determined by our sequencing data (**Fig 2C**). These findings increased confidence that the FACS sorting has captured a representative snapshot of the POMC neuronal population.

**Figure 6.**
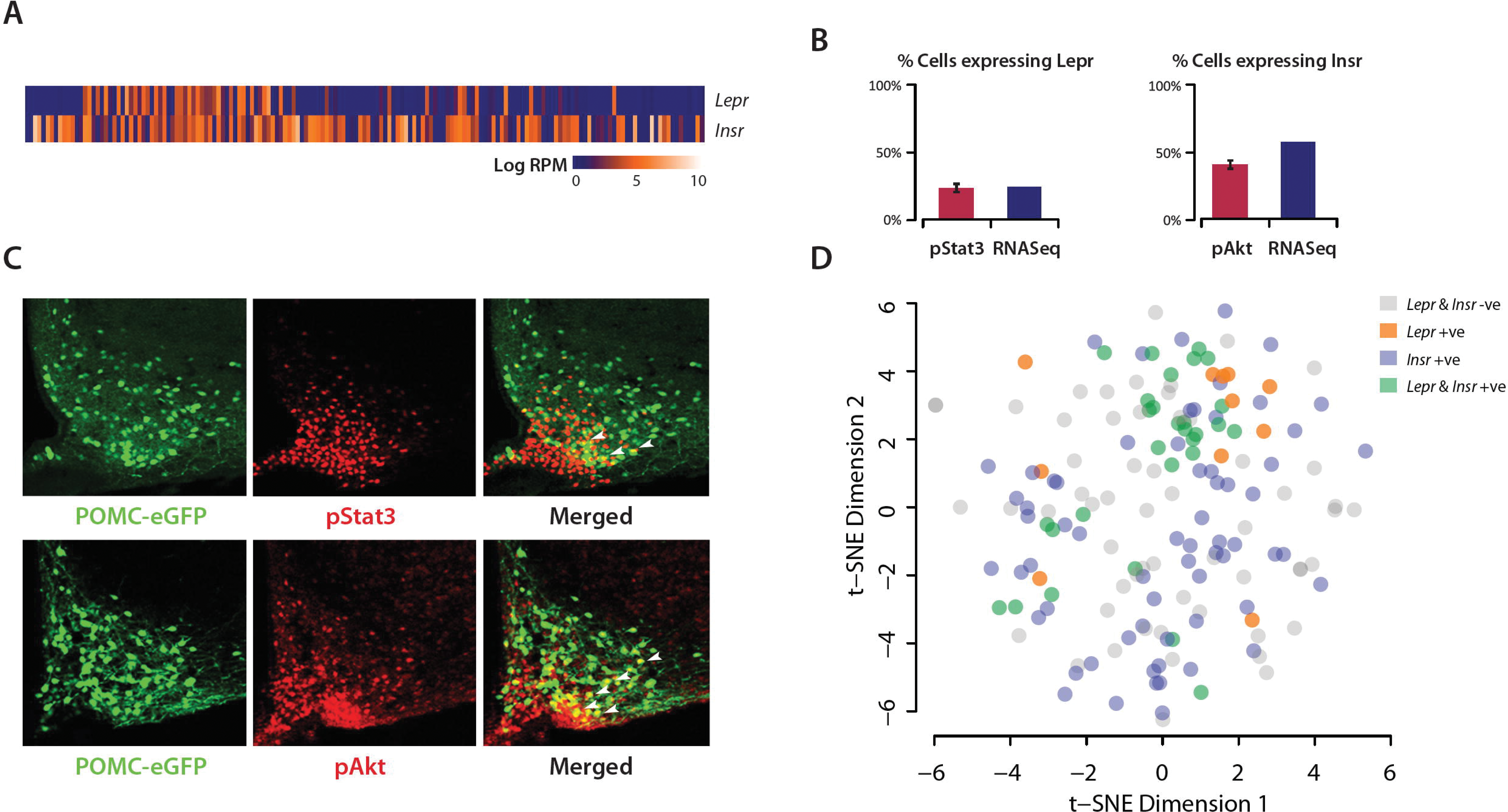
*Lepr* and *Insr* expression in hypothalamic POMC neurons. **(A)** We detected 30 *Lepr* and 83 *Insr* expressing cells out of the 163 cells in our sequencing cohort. **(B)** The percentages were compared to the number of cells that exhibited pStat3 and pAkt immunoreactivity post-administration of leptin and insulin (respectively). **(C)** Representative images showing POMC-eGFP, immunoreactivity of pStat3 (upper panel) and pAkt (lower panel). Cells that were double-stained were marked with white arrows. **(D)** The t-SNE plot shows a lack of clear clustering for *Lepr* and *Insr* expressing cells, indicating the expressions of the 2 receptors alone were not sufficient to drive segregation at the whole transcriptome level.

## Discussion

We describe here the use of single cell RNA sequencing to determine the transcriptomes of 163 individual murine POMC neurons. Unexpectedly, 25% of POMC positive neurons also express high levels of *Agrp*. The transcriptomes of these neurons are comparable with those of previously published pooled AgRP neurons (Henry et al., 2015), indicating that these are not an artefact but likely to be a real subpopulation of canonical AgRP neurons. Additionally, when we examine the raw data from Henry et al, we find that AgRP/NPY GFP neurons also express low levels of POMC (Henry et al., 2015). This phenomenon likely reflects the shared developmental origins of POMC and AgRP cells (Padilla et al., 2010). POMC has been shown to be broadly expressed in hypothalamic progenitors, with up to 25% of the mature AgRP/NPY population sharing a common progenitor with POMC neurons (Padilla et al., 2010). The A^+^P_+_ population we describe here could be cells where POMC transcription is never fully silenced, even after the differentiation into AgRP/NPY neurons. Another possibility is that the high *Agrp/Npy* expressing neurons could be newly committed AgRP/NPY neurons from these progenitor cells, where *Pomc* expression is yet to be fully suppressed. Regardless of the explanation, given that Pomc and AgRP genetic tools are in wide use, it is important for the field to know that a ‘*Pomc*-specific’ deletion of a given gene would also be removing the gene from a significant proportion of AgRP neurons. It is possible for example, the mild phenotype seen in the mice where *Lepr* is ‘specifically’ deleted from POMC neurons (Balthasar et al., 2004) could be in part due to *Lepr* deletion from a proportion of AgRP neurons.

The remaining 118 P^+^ neurons display striking heterogeneity and fall into four clusters. A few important points to keep in mind while interpreting these data. First, these groups have emerged in an unbiased fashion after achieving a certain statistical threshold, which means that they are not invariant. Thus increasing the threshold would invariably result in more clusters. Second, these groupings are not solely characterised by small numbers of unique genes; rather they are driven by a ‘signature’ expression profile of a whole repertoire of genes. Thus while the driver genes (see **Figure 5**) are on average the highest expressed in their particular group, many transcripts are not unique to a single group and are not necessarily expressed in all of the neurons in any given group. Third, these different clusters are likely to be dynamic, depending on the nutritional state of the animal at the time the brain was obtained, with the mice in this particular experiment being ad libitum fed.

The group with the highest expression of *Pomc* and *Cartpt* is cluster 4, and based on its expression profile of GPCRs, appears to be responsive to corticotrophin releasing hormone, hypocretin, prolactin, Npy, CCK and neuromedin U. The largest group is cluster 1, which based on its repertoire of GPCR expression, responds to vasopressin, melanocortin peptides (via the MC3R), serotonin and somatostatin. Cluster 1 is notable in that, compared to the other 3 groups, it expresses the highest levels of a wide variety of plasma membrane ion channels. We acknowledge that further experiments will be required to functionally characterize these neurons. However, we speculate that the enrichment of distinct classes of surface receptors on the different neuronal clusters possibly speaks to the temporal nature of the downstream effector function, with signalling through ion channels typically bringing about rapid changes, and circulating humoural factors providing longer term signals.

Two drugs that are used for weight-loss currently on the market are the serotonin receptor agonist Locaserin (Smith et al., 2010) and the GLP1 analogue Liraglutide (Astrup et al., 2009). There is evidence that both mediate their effects on food intake via ARC POMC neurons expressing 5HT_2C_R (Burke et al., 2016; Heisler et al., 2002) and GLP1R (Secher et al., 2014), respectively. Our data provides two additional pieces of information (**Figure 3**). First, that less than 10% (11/118) of ARC POMC neurons express the GLP1R. Second, of the POMC neurons expressing either 5HT_2C_R and GLP1R, only 3 neurons express both receptors. Thus it is likely that different subgroups of POMC neurons mediate the satiety effects of each compound, and poses the question of whether a potential combinatorial therapy, without increasing individual drug-dose, might possibly result in increased weight-loss.

As previously mentioned, the 45 A^+^P_+_ neurons are very distinct from the 118 P^+^ neurons, clearly segregating into a separate group. This is particularly true when one examines the expression of the leptin and insulin receptors in these groups. The low percentage (12%) of canonical POMC neurons that express the leptin receptor is consistent with the actions of leptin on these neurons being largely indirect (Cowley et al., 2001; Vong et al., 2011). Although we did not specifically or comprehensively evaluate an *Agrp* positive population of neurons, the much higher percentage of LepR positivity in the cells where we documented high levels of *Agrp* expression (57%) support the idea that AgRP neurons are more likely to be direct targets of leptin (Cowley et al., 2001). Our data also provide some evidence of autocrine/paracrine regulation by POMC neurons themselves, with 10% of P^+^ neurons expressing *Mc4r* and nearly 40% expressing the *Mc3r*. One could speculate that there may be amplification of the melanocortin system, whereby P^+^ neurons that respond directly to leptin go on to signal to *Mc3r/Mc4r* expressing neurons, but this remains to be determined.

In contrast, the expression of insulin receptors is comparable between the A^+^P_+_ and P^+^ neurons, indicating that insulins actions on both these populations of neurons are likely to be direct (Hill et al., 2010; Konner et al., 2007).

In conclusion, single cell RNAseq has revealed previously unappreciated heterogeneity in POMC neurons. We have provided a comprehensive atlas, which we hope will be of utility to basic and translational researchers attempting to better understand the fundamental nature of regulation of energy balance by the hypothalamus and to manipulate these systems for therapeutic benefit.

## Material and Methods

## Animals

7-9 week old male *Pomc-eGFP* transgenic mice (Cowley et al., 2001) (kind gift from Prof. Malcolm Low) were housed under a standard 12hr light-dark cycle and were fed *ad libitum*. All studies were approved by the local Ethics Committee and conducted according to the UK Home Office Animals (Scientific Procedures) Act 1986.

## Isolation of single hypothalamic POMC-eGFP neurons using Fluorescence-activated Cell Sorting (FACS)

*Ad libitum* fed *Pomc-eGFP* mice were sacrificed by cervical dislocation and tissue from the medial basal hypothalamus (MBH) region was micro-dissected into Hibernate A medium without calcium and magnesium (BrainBits, Springfield, IL, USA). The tissue was then digested with Papain (20U/ml, Worthington, Lakewood, NJ, USA) for 30min at 37 °C with mild agitation, followed by trituration in Hibernate A medium containing DNase I (0.005%, Worthington). The cell suspension was then filtered through a 40 μm cell strainer into a new falcon tube.

Fluorescence-activating cell sorting was performed using an Influx Cell Sorter (BD Biosciences, San Jose, CA, USA) utilising a protocol that was previously used for single-cell isolation (Macaulay et al., 2016; Schulte et al., 2015). The cell gating was set according to cell size (FSC), cell granularity (SSC), FSC pulse-width for singlets, fluorescence at 488nm/532nm for GFP and 647/670nm for nuclear stain with DraQ5 (Biostatus, Shepshed, Leicester, UK). Cells were sorted direct into a 96 well plate containing lysis buffer with RNase inhibitor.

## Whole-transcriptome amplification and single-cell RNA sequencing

RNA from the single cell lysate was reverse transcribed and PCR amplified (19 cycles) into double stranded cDNA using Clontech SMART-Seq v4 Ultra Low Input RNA Kit (Takara Clontech, Mountain View, CA, USA). 150pg of amplified cDNA was then used to generate barcoded sequencing libraries using Illumina Nextera XT library preparation kit (San Diego, CA, USA) according to manufacturer’s manual. Samples failing to show amplification were removed from the experiment.

The sequencing libraries were normalized to 10nM concentration and combined into pools of 96. The pooled libraries were sequenced on 4 lanes of an Illumina HiSeq 4000 instrument at single-end 50bp (SE50), yielding an average of 15.7 million reads per cell. Library preparation was performed by the Genomics and Transcriptomic Core at the Institute of Metabolic Science. The sequencing was performed at the Genomics Core, Cancer Research UK Cambridge Institute. All the raw data in this manuscript has been submitted to GEO, accession number GSE92707.

## Leptin and Insulin administrations

For pSTAT3 immuno-labelling, mice were fasted overnight and leptin (3mg/kg, PeproTech, Rocky Hill, NJ, USA) or saline was injected intraperitoneally. 45 minutes after leptin administration, the animals were then sacrificed by a lethal overdose of pentobarbital (60mg/kg) then perfused transcardially with PBS followed by 4% paraformaldehyde. For pAKT immunofluorescence overnight fasted animals were administered with Actrapid (2mU/kg, Novo Nordisk, Bagsværd, Denmark) intravenously. Animals were sacrificed 15 minutes after administration and perfused as described before.

## Immunohistochemistry for POMC-eGFP, pStat3 and pAkt

Brains were collected immediately after perfusions and fixed for 2 further hours in a 15% sucrose- 4% paraformaldehyde solution. The brains were cryo-preserved in 30% sucrose PBS overnight and then embedded in OCT (Bright, Huntingdon, UK) and stored at −80°C until further processing. Frozen coronal sections were sliced at 30 µm and processed for immunofluoresence. For pSTAT3 immunostaining, sections were pre-treated for 20 min in 1% NaOH and then placed overnight with a rabbit anti-pSTAT3 antibody (1:200, Tyr-705 Cell Signalling). For pAKT immunostaining, sections were incubated with anti-pAKT antibody (1:100, Ser-473 Cell Signalling) overnight. The primary antibodies were localized with Dylight 594 Goat anti-Rabbit IgGs (Vector; 1:500). Coronal images of Arcuate nucleus were taken using a Laser-scanning confocal microscope (LSM 510, Zeiss, Oberkochen, Germany) and eGFP positive cells and double labelled GFP-pSTAT3- or GFP-pAKT immune-reactive cells were counted using Image J.

## Immunohistochemistry for Mc4r-MAPT, and Pomc

B6.Cg-Tg(Mc4r-MAPT/Sapphire)21Rck/J male mice were perfused transcardially via a 23 gauge needle placed in the left ventricle with 100 ml of 0.1M heparinized PBS, pH 7.4, followed by 100 ml of 4% paraformaldehyde in PBS, and the fixed brains were cryoprotected in 30% sucrose for 48h at 4C. Coronal hypothalamic sections of 30 μm thickness were prepared on a freezing microtome. Free floating sections were incubated for 15 min in 1% hydrogen peroxide, washed 2 times in PBS, blocked 2h in 0.3% Triton X-100 and 5% NGS in PBS, before being incubated overnight at 4° C in rabbit anti-proopiomelanocortin precursor (1:200, Phoenix Pharmaceuticals) in 0.3% Triton X-100 and 5% NGS. Sections were then washed and incubated for 2 h with Alexa Fluor 594 goat anti-rabbit IgG. Sections were washed and incubated overnight at 4° C in chicken anti-GFP (1:500, Abcam). Sections were washed and incubated in Alex Fluor 488 goat anti-chicken IgG for 2h. Sections were floated onto superfrost Plus microscope slides (Fisher), and coverslipped with Vectashield (Vector). Tissue sections were digitized using a Zeiss Axioplan fluorescent microscope, and areas of interest were outlined based on cellular morphology.

## Bioinformatics

The quality of sequence reads was examined using FastQC and Multiple Genome alignment (MGA) was used to rule out contamination from other DNA sources throughout the wet experimental procedures. Reads were aligned to the mouse genome (GRCm38, Ensembl v74) using Tophat version 2.0.11 (Kim et al., 2013) and the raw counts at the gene level were determined using Cufflinks version 2.2.1 (Trapnell et al., 2010). The above analyses were performed at the High-performance computing cluster at the Research Computing Service, University of Cambridge.

The raw counts from cells were normalised and batch correction was performed using edgeR (Robinson et al., 2010) and limma (Ritchie et al., 2015) packages. Rtsne package was used to generate t-SNE plots (Maaten and Hinton, 2008) and SC3 package was used to perform the clustering analysis (Kiselev et al., 2016). GeneSpring GX software (Agilent) was used to generate the heatmaps. Gene abundance is represented in log RPM, which stands for the base-2 logarithm of Reads per Million. The plots were generated using ggplot2 package, Microsoft Excel and GeneSpring GX. Gene annotations and pathway analysis were perform using Ingenuity Pathway Analysis (Qiagen, Redwood City, CA, USA)

## Acknowledgements

This work was supported by the UK Medical Research Council (MRC) Metabolic Disease Unit (MRC_MC_UU_12012/1 & MRC_MC_UU_12012/5), a Wellcome Trust Strategic Award (100574/Z/12/Z) and the Helmholtz Alliance ICEMED.

